# Differential activation of parathyroid and renal Ca^2+^-sensing receptors underlies the renal phenotype in autosomal dominant hypocalcemia 1

**DOI:** 10.1101/2021.02.01.429170

**Authors:** Wouter H. van Megen, Rebecca Tan, R. Todd Alexander, Henrik Dimke

## Abstract

**Background:** Parathyroid Ca^2+^-sensing receptor (CaSR) activation inhibits parathyroid hormone (PTH) release, while activation of CaSRs in kidneys and intestine attenuates local transepithelial Ca^2+^ transport. In patients with autosomal dominant hypocalcemia 1 (ADH1) due to activating *CASR* mutations, treatment of symptomatic hypocalcemia can be complicated by treatment-induced hypercalciuria, resulting in nephrocalcinosis and renal insufficiency. Although CaSRs throughout the body are activated by increased extracellular Ca^2+^ concentrations, it is not understood why some ADH1 patients have reduced PTH, but not hypercalciuria at presentation, despite CaSR expression in both kidney and parathyroid.

**Methods:** Activation of the CaSR was studied in mouse models and a ADH1 patient. *In vitro* CaSR activation was studied in HEK293 cells.

**Results:** Mice with a gain-of-function mutation in *Casr* are hypocalcemic with reduced plasma PTH levels. However, renal CaSRs are not activated as indicated by normal urinary calcium handling and unaltered renal *Cldn14* expression. Activation of renal CaSRs only occurred after increasing plasma Ca^2+^ levels. Similarly, calcimimetic administration to wildtype mice induced hypocalcemia without activating renal CaSRs. Moreover, significant hypercalciuria was not observed in an ADH1 patient until blood Ca^2+^ was normalized. *In vitro* experiments suggest that increased CaSR levels in the parathyroid relative to the kidney contribute tissue-specific CaSR activation thresholds.

**Conclusion:** Here we delineate tissue-specific CaSR activation thresholds, where parathyroid CaSR overactivity can reduce plasma Ca^2+^ to levels insufficient to activate renal CaSRs, even with overactivating mutations. These findings may aid in the management of ADH1 patients.

## Introduction

Plasma ionized calcium (Ca^2+^) concentrations are kept within a narrow concentration range of 1.09–1.35 mM (1). Ca^2+^ activates the central Ca^2+^-sensing receptor (CaSR) in the parathyroid, thereby reducing the release of parathyroid hormone (PTH) to reduce plasma Ca^2+^ concentrations (2–4). PTH is a key stimulator of Ca^2+^ reabsorption in the kidney and Ca^2+^ mobilization from bone, while it also indirectly regulates intestinal Ca^2+^ absorption by stimulating the formation of the active vitamin D metabolite 1,25(OH)_2_D_3_ (1).

In addition to PTH-dependent regulation of Ca^2+^ transport in bone, kidney and intestine, local activation of the CaSR in these peripheral tissues allows for regulation of Ca^2+^ transport through a direct response to changes in plasma Ca^2+^ levels (1, 5, 6). In the kidney, the CaSR is highly expressed on the basolateral membrane of the thick ascending limb of Henle’s loop (TAL) (6). Activation of the CaSR in this segment reduces paracellular Ca^2+^ reabsorption due to an increased expression of the pore-blocking claudin-14 (CLDN14), which is hardly expressed under basal conditions (6–8). In fact, CaSR activation in the kidney plays a crucial role in the defense against hypercalcemia (9). Furthermore, we have found that CaSR activation in the intestine decreases the expression of the epithelial Ca^2+^ channel transient receptor potential vanilloid 6 (TRPV6), thereby attenuating intestinal Ca^2+^ absorption (10). Thus, activation of the CaSR in the kidney and intestine limits Ca^2+^ transport in these tissues by regulating the expression of CaSR-sensitive genes. It is generally believed, but not known, whether these peripheral CaSRs in kidney and intestine are activated at the same plasma Ca^2+^ concentrations as the CaSR in the parathyroid

Gain-of-function mutations in *CASR* lead to autosomal dominant hypocalcemia type 1 (ADH1). ADH1 is characterized by hypocalcemia with inappropriately low plasma PTH concentrations, due to CaSR activation in the parathyroid (11, 12). Frank hypercalciuria is most frequently encountered when correcting the hypocalcemia and not at presentation (11–15), even though activating CaSR mutations are found in receptors expressed in both the parathyroid and the kidney. Importantly, since treatment-induced hypercalciuria increases the risk of nephrolithiasis, calcifications and ultimately chronic kidney disease in patients with ADH1, treatment is generally avoided in asymptomatic patients (11). It is not clear why ADH1 patients present with reduced PTH, but not hypercalciuria, with CaSR expression in both kidney and parathyroid.

We therefore studied the activation of the CaSR across different tissues in response to changes in plasma Ca^2+^ levels in a mouse model of ADH1 (Nuf) and after administration of a calcimimetic, etelcalcetide hydrochloride (16–18). The expression of *Cldn14* in the kidney was determined as an indicator of renal CaSR activation, based on previous studies showing its expression is highly regulated by CaSR activation (7, 8) and since CaSRs have been detailed to use different signaling pathways in kidney and parathyroid (19).

We show here that genetic and pharmacological CaSR activation induces hypocalcemia with low urinary Ca^2+^ excretion, while the observation that the expression of renal and intestinal CaSR-sensitive genes was unaltered is in line with peripheral receptors not being activated. However, when plasma Ca^2+^ levels were increased in Nuf mice, but still lower than wild-type animals, we observed inappropriately increased urinary Ca^2+^ excretion and increased renal *Cldn14* expression. Similar findings were observed in a patient suffering from ADH1. Overall, our findings suggest an intricate interplay between the CaSR in the parathyroid and peripheral CaSRs, with different activation thresholds, where renal CaSRs are not activated at baseline, because CaSR activity in the parathyroid reduces PTH secretion and plasma Ca^2+^ levels. These findings are relevant for the management of ADH1 patients with hypocalcemia, suggesting that plasma Ca^2+^ should be increased sufficiently to prevent symptoms of hypocalcemia, but not raised to the normal range in order to avoid hypercalciuria and related sequelae.

## Methods

### Animal experiments

Nuf mice (C3;102-Casr^Nuf^/H.), which harbor an activating mutation (L723Q) in the fourth transmembrane region of the CaSR, have been described previously and were obtained through MRC Harwell via the European Mouse Mutant Archive (18). All animal experiments were conducted in accordance with Danish Law under animal experimental permits #2014-15-0201-00043 and #2019-15-0201-01629.

### Experimental protocol 1

Experiments were performed on sex-matched and age-matched wild type (WT) (*n*=9 animals, average age 10 weeks), heterozygous (Nuf/+) (*n*=15 animals) and homozygous (Nuf/Nuf) (*n*=10 animals). All mice were randomly assigned to *i.p.* injections of either 7.5 mg/kg furosemide (Furix; Takeda Pharma A/S, Taastrup, Denmark) or vehicle (0.9% NaCl). Animal weights were 22.62 g ± 2.77 g (SD) for WT, 22.57 g ± 3.09 g for Nuf/+ and 22.89 g ± 3.11 g for Nuf/Nuf mice. Spot urine was collected at baseline and 2 hours after injection; furosemide-sensitive Ca^2+^ excretion was calculated by subtracting baseline Ca^2+^ excretion from urine Ca^2+^ excretion after furosemide injection. One-week post-furosemide injection (when the animals were aged 12 weeks on average), bone mineral content, bone area, and bone mineral density were determined using dual-energy X-ray absorptiometry (DEXA) by means of a PIXImus2 densitometer with software version 1.44 (Lunar Cooperation, Fitchburg, WI, USA). Subsequently, all animals were transferred to individual metabolic cages in a temperature-controlled (25 °C) environment on a 12-hour light/12-hour dark cycle. Animals had *ad libitum* access to a standard rodent maintenance diet containing 0.2% Mg^2+^, 0.7% Ca^2+^ and 600 IU vitamin D3 (TD.00374; Altromin Spezialfutter GmbH & Co. KG, Lage, Germany) and drinking water. After acclimatization for 3 days, 24h urine was collected. Animals were subsequently anesthetized with 1.5% isoflurane (Nicholas Piramal Ltd., London, UK) and blood was drawn from the vena cava. Mice were then euthanized by cervical dislocation and kidneys, duodenum and colon were isolated as reported before (20).

### Experimental protocol 2

8 WT and 8 Nuf/+ mice (age 16-17 weeks) were placed on a 2%-supplemented Ca^2+^ diet (Envigo, Indianapolis, IN, USA) for 7 days. First, they received the diet in their standard cages for 3 days and were subsequently transferred to individual metabolic cages, where they continued receiving the high Ca^2+^ diet. All animals had *ad libitum* access to food and water and were housed in a controlled environment, as described above. After 3 days of acclimatization, a 24h urine was collected. Mice were subsequently anesthetized and euthanized under 1.5% isoflurane as described above.

### Experimental protocol 3

8 WT and 7 Nuf/+ mice (age 16-17 weeks) were housed in individual metabolic cages as described above. After acclimatization for 3 days, baseline 24h urine was collected. All mice were then switched to a diet supplemented with dihydrotachysterol (DHT), a synthetic reduced vitamin D analog which is activated in the liver and is independent of renal hydroxylation (0.4% w/v dihydrotachysterol (Sigma-Aldrich, St. Louis, MO, USA) per 100 g of standard rodent food) for 3 days; 24h urine was collected for all days (21). Mice were anesthetized and euthanized as described above.

### Experimental protocol 4

5 WT and 7 Nuf/+ female mice (age 8-11 weeks) were used in the experiment. To increase plasma Ca^2+^ towards normal levels in Nuf mice, only Nuf/+ mice were placed on a lower dose DHT-enriched diet (0.04% w/v dihydrotachysterol (Sigma-Aldrich) per 100 g of standard rodent food) for 7 days, whereas WT mice received standard diet. Mice were first maintained in regular cages, then placed in individual metabolic cages for acclimatization for 3 days, after which 24h urine was collected on the 7^th^ day. Mice were anesthetized and euthanized as above.

### Experimental protocol 5

18 C57BL7/6J male mice (average age 10 weeks) were randomly divided into 3 groups with a similar average body weight. One group of animals served as controls, one group received infusion with 5 mg/ml etelcalcetide hydrochloride (Parsabiv; Amgen) via Alzet osmotic 2002 minipumps (Alzet, Cupertino, CA, USA) with a delivery rate of 0.5 μl/hr, and one group received infusion with 20 mg/ml etelcalcetide hydrochloride via Alzet osmotic 2001 minipumps (Durect Corporation, Cupertino, CA, USA) with a delivery rate of 1.0 μl/hr. Three days after insertion of minipumps, animals were housed in individual metabolic cages. After acclimatization for 3 days, baseline 24h urine was collected on day 7. Mice were anesthetized and euthanized as above.

### Biochemical measurements

Blood ionized Ca^2+^ concentrations were measured by means of an ABL835 FLEX analyzer (Radiometer, Copenhagen, Denmark). Plasma was isolated from the remaining blood by centrifuging at 4500 *g* for 10 minutes at 4 °C and subsequently stored at −80 °C for further analyses. Urine Ca^2+^ concentrations were measured using a colorimetric assay (22). Plasma and urine creatinine concentrations were measured using the Jaffé method with an ABX Pentra Creatinine 120 CP kit (HORIBA ABX SAS, Montpellier, France) on a Microlab 300 analyzer (ELITechGroup, Puteaux, France), according to the manufacturer’s protocol. For experimental protocol 4 and 5, plasma and urine creatinine concentrations were measured by the Department of Clinical Biochemistry and Pharmacology of Odense University Hospital. Intact plasma parathyroid hormone (PTH) levels were determined with a mouse PTH 1-84 ELISA kit (Immutopics International, San Clemente, CA) as before (7).

Fractional excretion of Ca^2+^ was calculated as

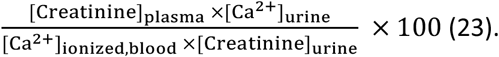

### RNA extraction and semi-quantitative PCR

Kidneys, duodenum and colon were snap-frozen in liquid nitrogen after isolation and stored at −80 °C for future RNA extraction. RNA was extracted from the tissues using TRIzol (Invitrogen, Carlsbad, CA, USA) and processed as previously (20, 24). In brief, after DNAse treatment (Invitrogen, Carlsbad, CA, USA), cDNA synthesis was performed using the SensiFAST cDNA Synthesis Kit (Bioline, London, UK). Then, qPCR was performed in triplicate for each sample on the QuantStudio 6 Pro Real-Time PCR System (Applied Biosystems, Waltham, MA, USA) using TaqMan Master Mix (Applied Biosystems, Waltham, MA, USA) and specific probes and primers (Table S1). Probes and primers were obtained from Applied Biosystems, Foster City, CA. For gene expression in the experiments regarding the Nuf mice, analysis was performed using the 2^−ΔΔCt^ approach with 18S rRNA as a reference gene. For the experiments with etelcalcetide hydrochloride, *Gapdh* was used as a reference gene. This was because *18S* showed significant differences between the groups.

### Immunohistochemistry

Immunohistochemistry was performed as described before (23). Kidneys were immersion-fixed in 10% formalin overnight at 4 °C. They were then stored in PBS until paraffin embedding. Embedding was performed using a Tissue-Tec Vacuum Infiltration Processor 6 (Sakura Finetek, Torrance, CA, USA) and then kidneys were cut into 2 μm thick sections with a HM 355S Automatic Microtome (Thermo Scientific, Kalamazoo, MI, USA). For immunostaining, sections were processed with Tissue-Tek Tissue-clear (Sakura Finetek, Brøndby, Denmark) and a graded series of ethanol. Subsequently, they were boiled for 10 minutes in Tris-EGTA buffer (10 mM Tris, 0.5 mM EGTA, pH = 9.0). Endogenous peroxidase activity and free aldehyde groups were blocked by incubating in 0.6% H_2_O_2_ and 50 mM NH_4_Cl in PBS. Then, sections were incubated overnight at 4 °C using a mouse monoclonal antibody against CLDN14 (Clone 5) produced in-house and characterized in another manuscript (manuscript attached). After washing, sections were incubated with secondary HRP coupled antibodies followed by addition of DAB^+^ Substrate Chromogen System (Dako Omnis, Glostrup, Denmark). For each experiment, 6 mice were tested per genotype and representative images are shown. For the immunohistochemical analyses of the parathyroid and renal cortex, sections were processed as described above and stained using a rabbit polyclonal antibody against the CaSR (PB9924, Boster Biological Technology, Pleasanton, CA, USA).

### Clinical data from ADH1 patient

Blood ionized Ca^2+^ and creatinine, urine Ca^2+^ and creatinine, and blood PTH were determined in the University of Alberta Hospital central laboratory by validated clinical biochemical protocols. Ionized blood Ca^2+^ concentrations between 1.09 and 1.25 mM are considered to be normocalcemia, while hypercalciuria is considered to be a urinary Ca^2+^/Crea ratio > 0.6. We included data that had either paired blood ionized Ca^2+^ and parathyroid levels or blood ionized Ca^2+^ and urine Ca^2+^ and creatinine levels. Due to the many samples collected over the life of this child, we averaged ionized Ca^2+^ and FECa or urine Ca^2+^/Crea over a range of ionized Ca^2+^ and plotted them in Figure 7. In consultation with the REB at the University of Alberta, formal ethical approval was not considered necessary, however, assent from the patient was provided as was formal written consent to include the clinical details of our subject by his guardian.

### *Cldn14* reporter assay in HEK293 cells

Cloning of the *Cldn14* promoter, which contains a CaSR responsive element is described separately (manuscript attached). HEK293 cells were purchased from the ATCC (Rockville, MD, USA) and culture in DMEM with 10% FBS and 5% penicillin, streptomycin and glutamine as previously (25). Cells were transfected via the calcium-phosphate method with the reporter construct, the pRL-TK renilla luciferase vector as an internal control and varying concentration of a CASR containing a myc tag vector (Origene Inc.). After 24 hr incubation with standard medium containing 1.5 mM calcium the cells were assayed for Cldn14 reporter expression using a luciferase assay system from Pormega Corp, according to the manufactures directions. Luciferase activity was assayed with 10 mL of extract on a DLR Ready, TD-20/20 Luminometer (Turner Design).

### Western blotting

Lysate from the luciferase assay was subjected to SDS Page electrophoresis and immunoblotting after mixing 3:1 with RIPA buffer as previously (7). A mouse primary anti-CaSR monoclonal antibody (1:2000, Gentex, Zeeland, MI) or a mouse primary anti-Myc (9B11) monoclonal antibody (1:1000, Cell Signaling Technology) was applied followed by a secondary horseradish peroxidase-coupled secondary antibody (1:5000, Santa Cruz Biotechnology, Santa Cruz, CA). This enabled detection of the CASR at the appropriate molecular weight after incubation with Western Lightning Plus ECL reagents (PerkinElmer, Boston, MA) and visualized using a Kodak Image Station 440CF (Kodak, Rochester, NY).

### Statistical analyses

Statistical analyses were performed using GraphPad Prism 8.3.0 (GraphPad software, San Diego, CA, USA). Comparisons between more than two groups were performed using one-way ANOVA with Tukey’s *post hoc* test or repeated measurements two-way ANOVA with Bonferroni’s *post hoc* test, as indicated in the figure legends. Analyses between two groups were performed using Student’s independent t-test (two-tailed). For all analyses, a *P* value < 0.05 was considered statistically significant. Data are presented as mean ± SEM

## Results

### Nuf mice are hypocalcemic but without renal Ca^2+^ wasting at baseline

As reported before, Nuf/+ and Nuf/Nuf mice are hypocalcemic, with Nuf/Nuf mice exhibiting more severe hypocalcemia than Nuf/+ mice (Figure 1A) (18). In addition, plasma PTH concentrations were reduced in Nuf/+ and Nuf/Nuf mice compared to WT mice (Figure 1B). Nuf/+ and Nuf/Nuf mice both had reduced urinary Ca^2+^ to creatinine ratio relative to wild-type mice, although the reduction was not significantly different between Nuf/+ and Nuf/Nuf mice (Figure 1C). Despite the reduced urinary Ca^2+^ to creatinine ratio observed in Nuf/+ and Nuf/Nuf mice, the fractional urinary excretion of Ca^2+^ was unaltered between all genotypes (Figure 1D), consistent with unaltered renal Ca^2+^ reabsorption. To investigate whether overactivation of the CaSR in the TAL alters paracellular Ca^2+^ reabsorption, which could be compensated distally, animals were challenged with furosemide. Furosemide inhibits sodium-potassium-chloride cotransporter 2 (NKCC2), thereby indirectly reducing Ca^2+^ reabsorption in the TAL (26). After furosemide injection, Ca^2+^ excretion was significantly increased in all genotypes (Figure 1E). However, furosemide-induced calciuresis was not significantly different in Nuf/+ or Nuf/Nuf mice compared to WT mice (Figure 1F). Together, these data suggest that although Nuf mice are hypocalcemic due to activation of the CaSR in the parathyroid and resultant PTH release, renal Ca^2+^ handling was unaltered.

**Figure 1.**
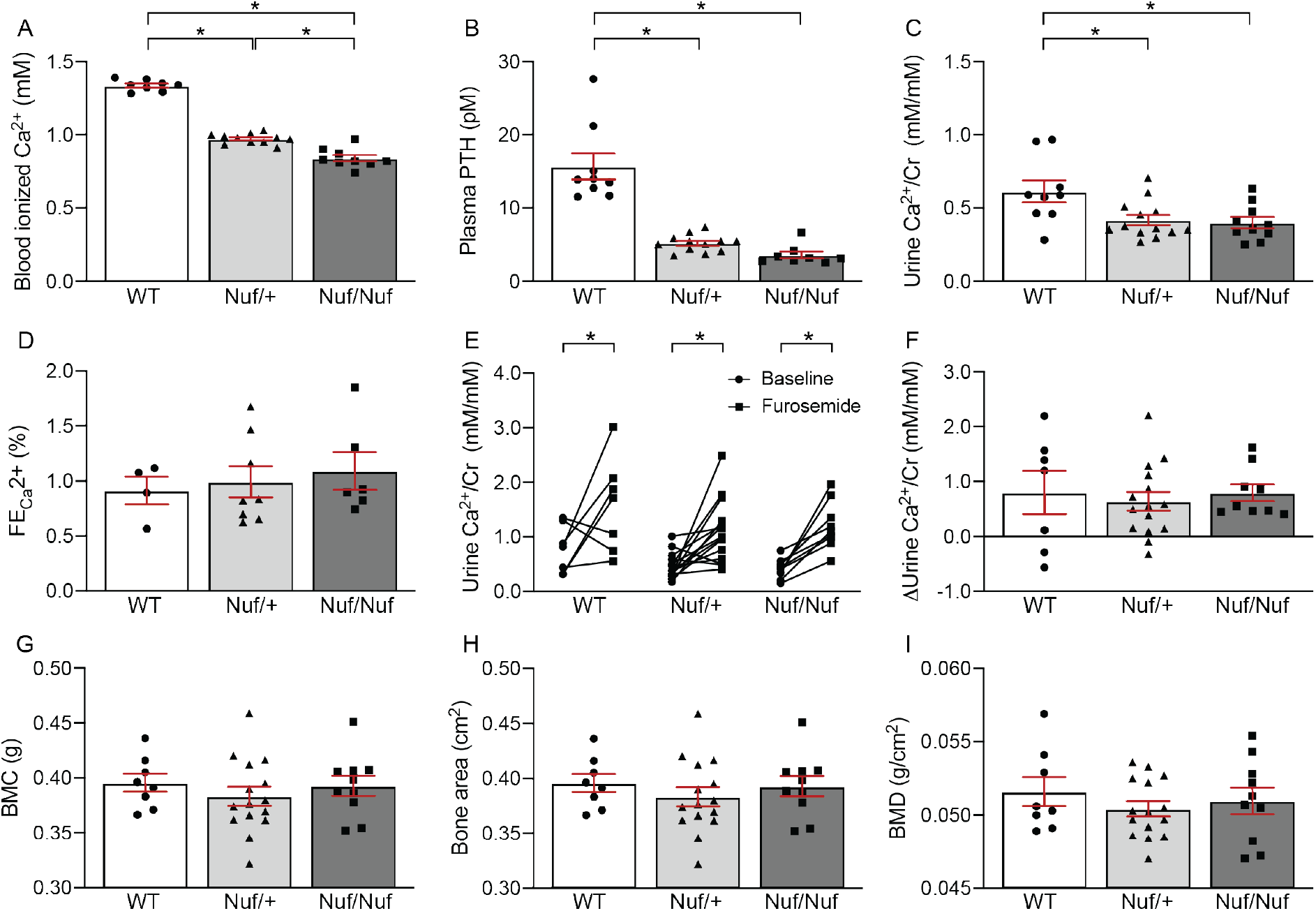
Ca^2+^ homeostasis in Nuf mice. Calcium levels in blood were significantly lower in Nuf/+ and Nuf/Nuf mice compared to WT animals. Ionized blood Ca^2+^ was also significantly lower in Nuf/Nuf mice compared to Nuf/+ mice (*n*=8-11) (A). Plasma intact PTH concentrations were significantly lower in Nuf/+ and Nuf/Nuf mice than in WT mice (*n*=9-12) (B). Urinary Ca^2+^ excretion was significantly lower in Nuf/+ mice and Nuf/Nuf mice compared to WT animals, but no significant differences were detected between Nuf/+ and Nuf/Nuf mice (*n*=9-13) (C). Fractional urinary excretion of Ca^2+^ (FECa^2+^) was similar in all genotypes (*n* = 4-8) (D). Administration of the diuretic furosemide significantly increased urinary Ca^2+^ excretion 2 hours after administration in all genotypes (*n*=7-15) (E). The furosemide-sensitive Ca^2+^ excretion (ΔCa^2+^/Cr) was not different between any of the genotypes (*n*=7-15) (F). Whole-body bone mineral content was not significantly different between Nuf mice and wildtype mice (*n*=8-15) (G). Total bone area was also unchanged (*n*=8-15) (H), while no differences were also present for whole-body bone mineral density (*n*=8-15) (I). Data are shown as mean ± SEM. *\*P* < 0.05 by one-way ANOVA with Tukey’s *post hoc* test (A-C) or two-way ANOVA (repeated measures) with Bonferroni’s *post hoc* test (E).

### Nuf mice have normal bone mineralization

Overactivation of CaSR in bone has been associated with bone abnormalities, including increased or reduced bone mineral density, reduced bone turnover and reduced bone formation (27, 28). Therefore, we measured whole-body bone mineral parameters of WT, Nuf/+ and Nuf/Nuf mice by DEXA. However, we were unable to find significant differences between genotypes in bone mineral content (Figure 1G), in total area bone (Figure 1H), or bone mineral density (Figure 1I). This suggests that in the Nuf/+ and Nuf/Nuf mice, the CaSR in the bone is not inappropriately activated unlike the parathyroid

### Renal and intestinal CaSRs are not activated in Nuf mice

To investigate if the renal CaSR is activated in the Nuf mice, we measured the expression of *Cldn14*. However, *Cldn14* expression was unaltered in Nuf/+ and Nuf/Nuf mice compared to WT mice (Figure 2A). In addition, despite the hypocalcemia in Nuf mice and reduced plasma PTH levels, the renal expression of the 1,25(OH)_2_D_3_-inactivating enzyme *Cyp24a1* was unaltered (Figure 2B). The expression of *Cyp27b1*, the proximal tubular enzyme responsible for generating 1,25(OH)_2_D_3_, was also unchanged in Nuf mice (Figure 2C). Immunohistochemistry showed unaltered localization of CLDN14 to the tight junctions of Nuf mice compared to WT mice (Figure 2D). These data are consistent with the unaltered fractional excretion of Ca^2+^ observed and indicate that at the lower plasma Ca^2+^ levels in the Nuf/+ and Nuf/Nuf mice, activation of the renal CaSR does not occur. Similarly, we have shown that activation of the intestinal CaSR in the duodenum and colon attenuates transcellular Ca^2+^ absorption by inhibiting TRPV6 activity and *Trpv6* expression (10). We therefore looked at the expression of genes mediating transcellular calcium absorption in these tissues. We found unaltered expression of *Trpv6* in the duodenum and colon of Nuf/+ and Nuf/Nuf mice, consistent with the intestinal CaSR not being activated at baseline in these mice (Figure 2E,F). Moreover, the expression of other calciotropic genes in the duodenum (Figure 2E) and colon (Figure 2F) was unaltered in Nuf/+ and Nuf/Nuf mice, although colonic expression of S100g (encoding calbindin-D9k) was significantly reduced in Nuf/Nuf mice compared to WT mice (Figure 2F). Together, these data suggest that in Nuf mice, the CaSR is only active in the parathyroid and not in bone, kidney or intestine.

**Figure 2.**
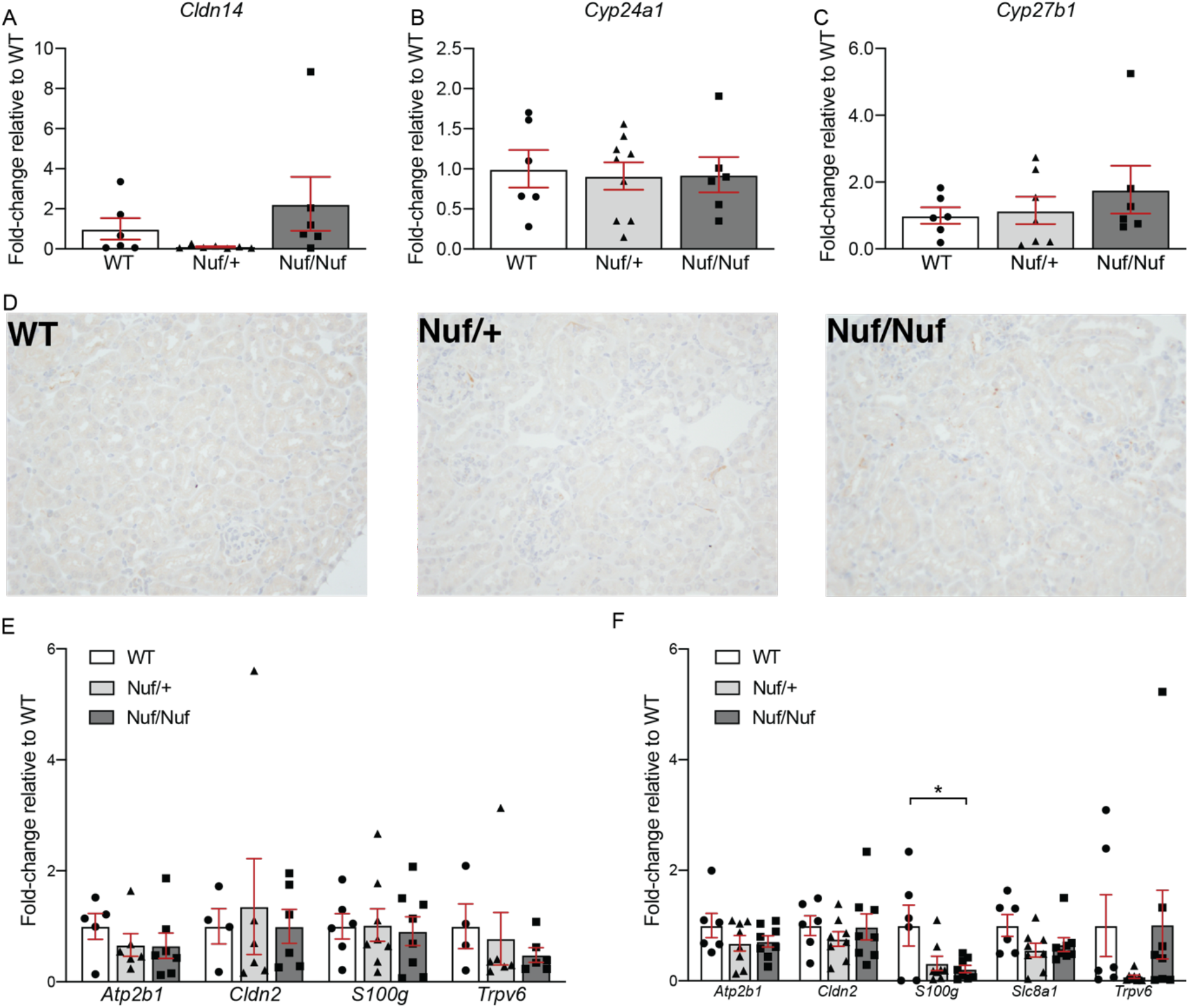
The renal and intestinal CaSR is not activated under baseline conditions in Nuf mice. The renal expression of *Cldn14*, a marker of CaSR activation in the kidney, was not significantly different between genotypes (*n*=6-7) (A). The renal expression of *Cyp24a1*, involved in the inactivation of 1,25(OH)_2_D_3_, was also unaltered in Nuf mice (*n*=6-9) (B). The renal expression of *Cyp27b1*, involved in the activation of 1,25(OH)2D3, was not statistically different between genotypes (*n*=6-7) (C). Localization of CLDN14 to the tight junctions of the TAL was not observed in any of the genotypes (D). The expression of genes involved in Ca^2+^ absorption in the duodenum, including *Atp2b1* (encoding PMCA1b), *Cldn2*, *S100g* (encoding calbindin-D9k) and *Trpv6* (a known CaSR-responsive gene) was not statistically different in Nuf/+ and Nuf/Nuf mice compared to WT mice (*n*=4-8) (E). The expression of *S100g* in the colon was significantly lower in Nuf/Nuf mice compared to WT mice, while the other colonic genes (including *Slc8a1*, which encodes NCX1) were not different between genotypes (*n*=6-8) (F). \**P* < 0.05 by one-way ANOVA with Tukey’s *post hoc* test. Data are shown as mean ± SEM.

### Dietary Ca^2+^ supplementation does not correct hypocalcemia or induce renal Ca^2+^ wasting in Nuf mice

Next, we administered a high (2% w/w) Ca^2+^ diet to WT and Nuf/+ mice for 6 days in an attempt to activate the renal CaSR in Nuf/+ mice (7, 8). Only heterozygous Nuf mice were used in this and the following experiments, since they reflect the heterozygous genotype of patients with ADH1 and showed similar renal Ca^2+^ handling as homozygous Nuf mice. However, Nuf/+ mice remained hypocalcemic (Figure 3A) with a low urinary Ca^2+^ to creatinine ratio (Figure 3B) and a similar fractional urinary excretion of Ca^2+^ as WT mice (Figure 3C). Moreover, dietary Ca^2+^ supplementation did not differentially activate the renal CaSR in Nuf/+ mice compared to WT mice, as indicated by the similar renal expression of *Cldn14* (Figure 3D). The expression of the vitamin D-converting enzymes *Cyp24a1* (Figure 3E) and *Cyp27b1* (Figure 3F) was also unaltered. Immunohistochemistry also revealed no changes in CLDN14 localization between Nuf/+ mice and WT mice (Figure 3G). No statistically significant differences were also found in calciotropic genes in the duodenum (Figure 3H) or colon (Figure 3I) of WT and Nuf/+ animals. These findings indicate that on a high Ca^2+^ diet, the renal and intestinal CaSRs are also not inappropriately activated in Nuf mice.

**Figure 3.**
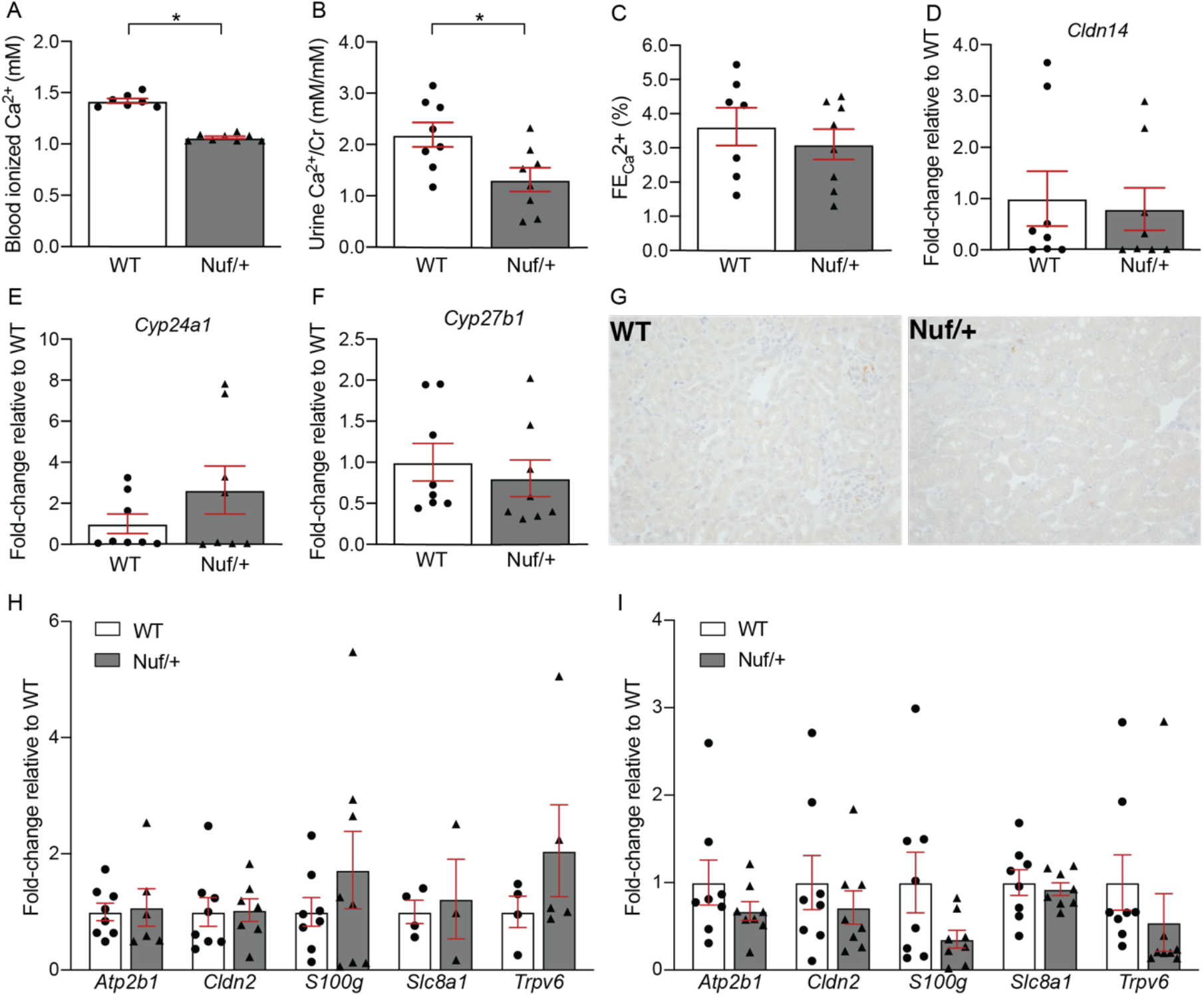
Ca^2+^ homeostasis in Nuf mice in response to dietary Ca^2+^ enrichment. After 6 days of a 2% Ca^2+^ diet, blood Ca^2+^ levels in Nuf/+ mice remained significantly lower than WT mice (*n*=7-8) (A). Urinary Ca^2+^ excretion was also significantly lower in Nuf/+ mice (*n*=8) (B). Fractional urinary excretion of Ca^2+^ (FE_Ca_^2+^ was not significantly different between both genotypes (*n*=7-8) (C). Renal *Cldn14* expression was not different bewtween Nuf/+ mice and WT mice on a high Ca^2+^ diet (*n*=8) (D). The renal expression of *Cyp24a1* and *Cyp27b1* was also not statistically different between Nuf/+ mice and WT mice (*n*=8) (E,F). For both genotypes, localization of CLDN14 to the tight junctions was not observed (G). A high Ca^2+^ diet did not differentially activate the intestinal CaSR in Nuf/+ mice, as indicated by the unaltered *Trpv6* expression (H,I). No significant differences were present for other genes involved in intestinal Ca^2+^ absorption in the duodenum (*n*=3-8) (H) or colon (*n*=8) (I). Data are shown as mean ± SEM. \**P* < 0.05 by Student’s independent t-test (two-tailed).

### Dietary DHT supplementation induces hypercalcemia and comparably activated the renal CaSR in Nuf/+ and WT mice

Active vitamin D regulates plasma Ca^2+^ by stimulating intestinal Ca^2+^ absorption (29). Therefore, the synthetic vitamin D analog DHT, which is activated in the liver, was added to the food of WT and Nuf/+ mice for 3 days to increase blood Ca^2+^ levels (21). This resulted in a similar blood Ca^2+^ concentration in Nuf/+ mice relative to WT mice, although both genotypes became hypercalcemic (Figure 4A). Plasma PTH concentrations remained slightly lower in Nuf/+ mice (Figure 4B). Treatment with DHT increased urinary Ca^2+^ excretion in both genotypes (Figure 4C). Remarkably, the increase in urinary Ca^2+^ excretion following DHT supplementation was significantly delayed in Nuf/+ mice compared to WT animals, i.e. was not observed until 3 days of treatment as opposed to the second day of treatment for WT animals (Figure 4C). Fractional urinary excretion of Ca^2+^ was not significantly different between genotypes (Figure 4D). Three days after dietary DHT supplementation, the expression of *Cldn14* was similar between Nuf/+ mice and WT mice (Figure 4E). In addition, the renal expression of *Cyp24a1* (Figure 4F) and *Cyp27b1* (Figure 4G) was not significantly different between the two genotypes. In contrast to the high Ca^2+^ diet, dietary DHT supplementation increased CLDN14 expression in both Nuf/+ and WT mice, with no obvious differences between the genotypes (Figure 4H). The expression of *Atp2b1* (encoding PMCA1b)*, Cldn2*, *S100g* and *Trpv6* in the duodenum was similar between both genotypes (Figure 4I), while the colonic expression of *Atp2b1*, *Cldn2*, *S100g*, *Slc8a1* (encoding NCX1) and *Trpv6* was also unaltered (Figure 4J). In parallel to the kidney, the expression of *Cyp24a1* was also not altered in the colon of Nuf/+ mice (Figure 4J).

**Figure 4.**
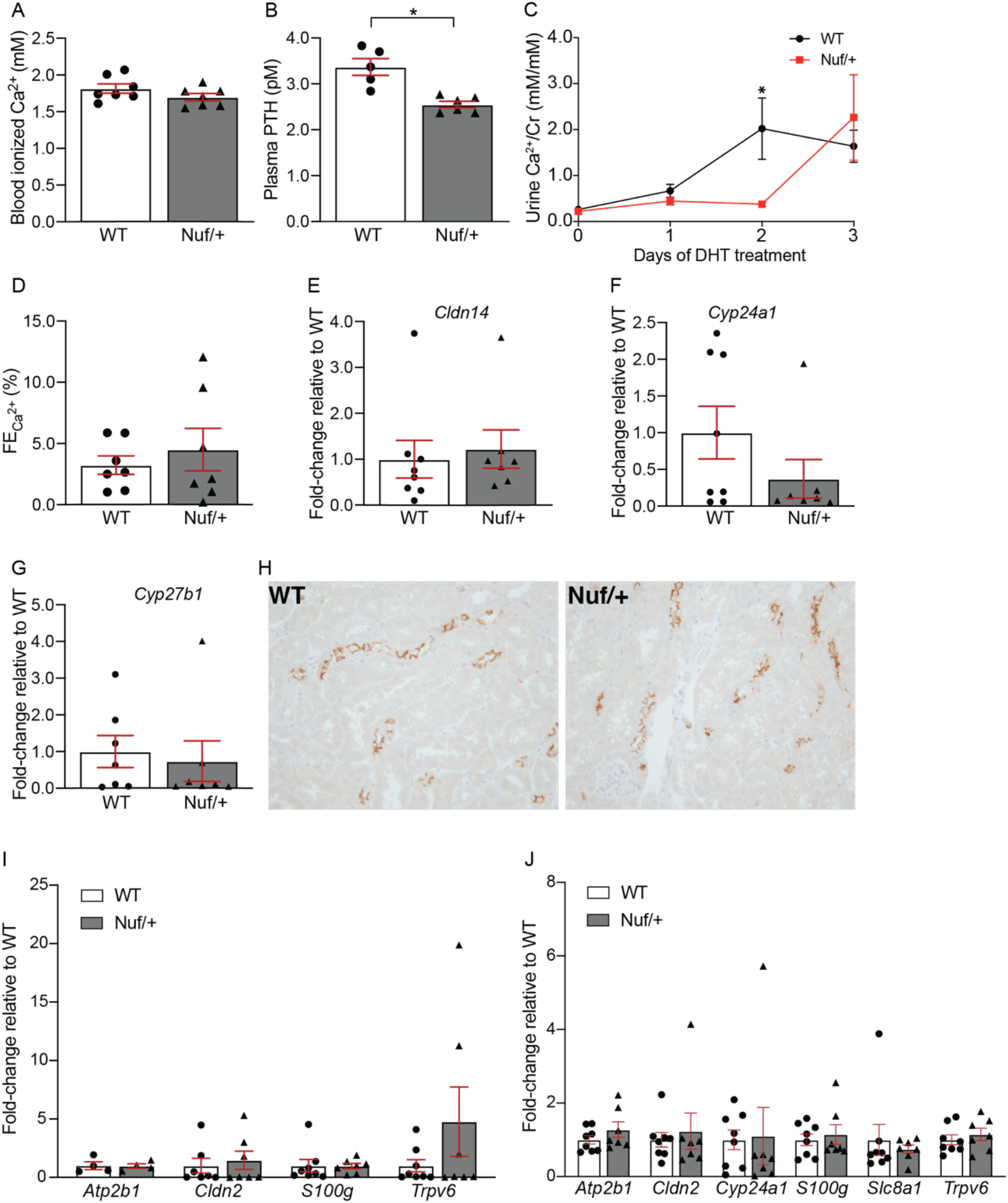
Effect of dietary DHT supplementation on Ca^2+^ balance. No significant changes in blood Ca^2+^ between WT and Nuf/+ mice were present after 3 days of dietary DHT supplementation (*n*=7) (A). Plasma PTH concentrations were significantly lower in Nuf/+ mice (*n*=5-6) (B). Although DHT increased urinary Ca^2+^ excretion in both genotypes, this increase was delayed significantly, by one day, in Nuf/+ mice compared to WT mice (*n*=7-8) (C). Fractional urinary excretion of Ca^2+^ (FE_Ca_^2+^) 3 days after DHT treatment was not significantly different between the two genotypes (*n*=7) (D). No differences in renal *Cldn14* expression were present between Nuf/+ mice and WT mice after 3 days of DHT supplementation (*n*=7-8) (E). The renal expression of *Cyp24a1* (*n*=7-8) and *Cyp27b1* (*n*=7) was also similar between Nuf/+ mice and WT mice (F,G). Immunohistochemistry showed a marked localization of CLDN14 to the tight junctions of Nuf/+ and WT animals treated with DHT for 3 days. No apparent differences were observed between Nuf/+ and WT animals (H). The expression of *Atp2b1*, *Cldn2*, *S100g* and *Trpv6* in the duodenum of Nuf/+ mice was not statistically different from WT mice (*n*=4-8) after dietary DHT administration (I). Similarly, DHT administration did not differentially affect the expression of *Atp2b1*, *Cldn2*, *Cyp24a1*, *S100g*, *Slc8a1* or *Trpv6* in the colon of Nuf/+ mice compared to WT mice (*n*=7-8) (J) \**P* < 0.05 for WT *vs.* Nuf/+ by Student’s independent t-test (two-tailed) (B) or by repeated measurements two-way ANOVA with Bonferroni’s *post hoc* test;, interaction between time and genotype was significant (*P* = 0.0352) (C). Data are shown as mean ± SEM.

### Differential activation of the renal CaSR after normalization of plasma Ca^2+^ levels in Nuf/+ mice

Since DHT treatment induced hypercalcemia in both genotypes (Figure 4A), we next attempted to raise plasma Ca^2+^ levels in Nuf/+ mice to levels comparable to steady-state Ca^2+^ levels in WT mice. Therefore, we used a lower dose of dietary DHT supplementation for 3 days and administered this only to Nuf/+ mice. After treatment with DHT, blood Ca^2+^ concentrations remained significantly lower in Nuf/+ mice than in untreated WT mice, although the difference between Nuf/+ mice and wild-type animals was smaller compared to previous experiments (1.34 mM (WT) *vs*. 0.97 mM (Nuf/+) under basal conditions (Figure 1A), compared to 1.34 mM (WT) *vs*. 1.19 mM (Nuf/+) (Figure 5A). Surprisingly, urinary Ca^2+^ excretion was significantly increased in Nuf/+ mice compared to WT mice (Fig 5B), while fractional urinary excretion of Ca^2+^ was also significantly higher in DHT-treated Nuf mice compared to WT mice (Figure 5C). A statistically significant increase in renal *Cldn14* expression was found in Nuf/+ mice compared to WT mice (Figure 5D). Furthermore, the expression of *Cyp24a1* was significantly higher in Nuf/+ mice than in WT mice (Figure 5E) and the expression of *Cyp27b1* was significantly decreased in Nuf/+ mice (Figure 5F). In addition to increased *Cldn14* expression in DHT-treated Nuf/+ mice, we also found CLDN14 expression increased in the TAL of Nuf/+ mice. No CLDN14 expression was detectable in the kidneys of untreated WT mice (Figure 5G). In the duodenum, the expression of *Slc8a1* was significantly increased compared to untreated WT mice (Figure 5H, while the colonic expression of *S100g* and *Trpv6* was significantly higher (Figure 5I). These results strongly suggest that the CaSR-CLDN14 pathway becomes activated in Nuf/+ mice after normalization of blood Ca^2+^ levels and is responsible for the increased urinary Ca^2+^ excretion. In addition, the significant increase in intestinal calciotropic gene expression likely facilitates increased dietary Ca^2+^ absorption in response to increased vitamin D.

**Figure 5.**
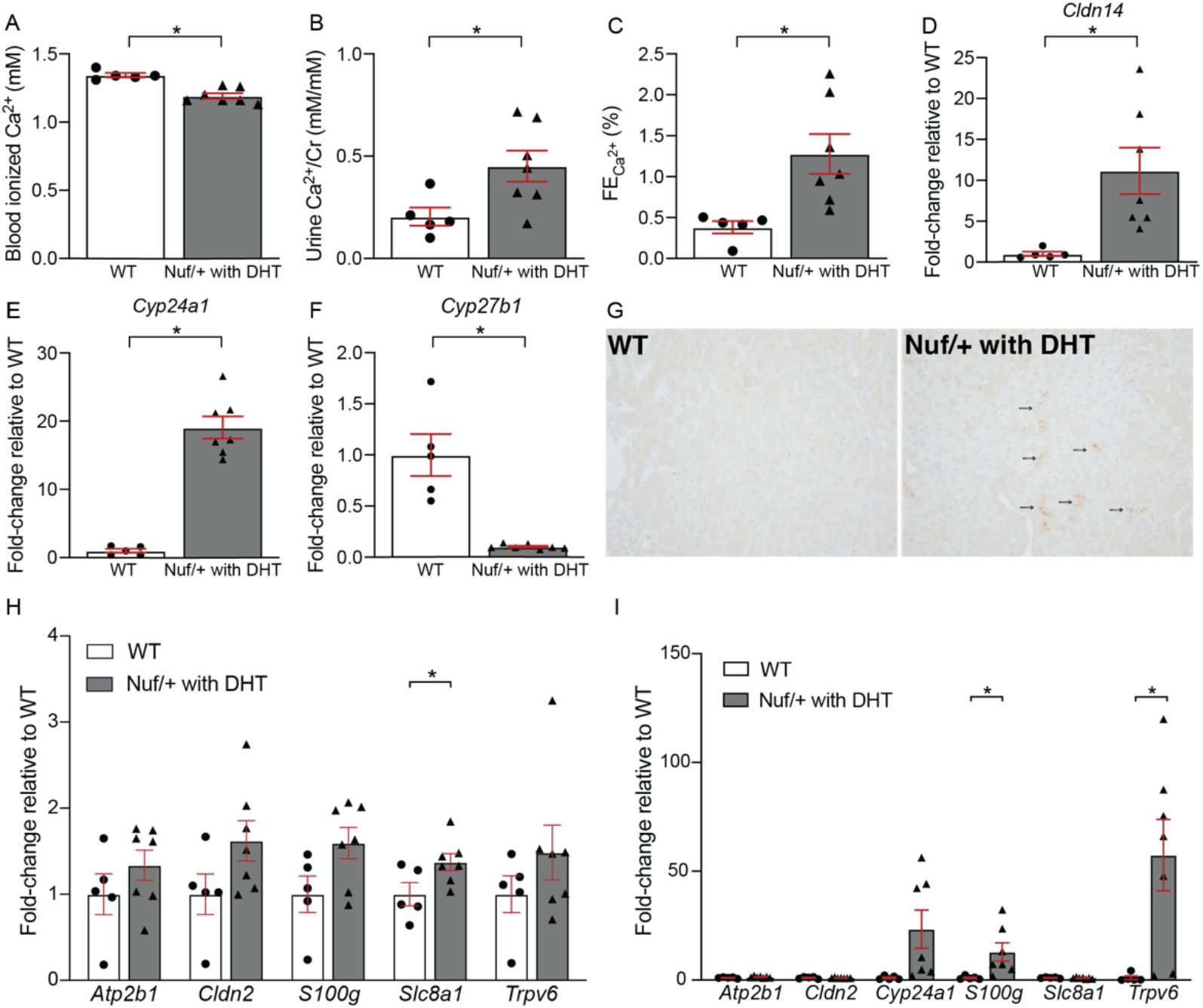
The renal CaSR becomes activated after 3-day DHT treatment in Nuf/+ mice. Plasma Ca^2+^ remained significantly lower in Nuf/+ mice treated with DHT for 3 days compared to untreated WT mice (*n*=5-7) (A). Urinary Ca^2+^ excretion was significantly higher in Nuf/+ mice following 3 days of DHT treatment compared to untreated WT mice (*n*=5-7) (B). Fractional urinary excretion of Ca^2+^ (FE_Ca_^2+^) was also significantly higher in DHT-treated Nuf mice than untreated WT mice (*n*=5-7) (C). The expression of *Cldn14* in the kidney was significantly higher in Nuf/+ mice compared to untreated WT animals (*n*=5-7) (D). In addition, increased renal *Cyp24a1* expression was present in Nuf/+ mice (*n*=5-7) (E). In contrast, the expression of *Cyp27b1* was significantly reduced in Nuf/+ mice (*n*=5-7) (F). Immunohistochemistry showed a marked localization of CLDN14 to the tight junctions of DHT-treated Nuf/+ mice, whereas this was absent-in WT animals (G). Dietary DHT administration to Nuf/+ mice increased the expression of *Slc8a1* in the duodenum, compared to untreated WT mice (*n*=5-7) (H). In the colon, DHT-treated Nuf mice exhibited an increased expression of *S100g* and *Trpv6* (*n*=5-7) (I). Data are shown as mean ± SEM. \**P* < 0.05 by Student’s independent t-test (two-tailed).

### Infusion of etelcalcetide hydrochloride leads to hypocalcemia without activating the renal CaSR

The calcimimetic etelcalcetide hydrochloride is a novel therapeutic for secondary hyperparathyroidism by activating the CaSR, thereby decreasing plasma Ca^2+^ and PTH concentrations (16, 17). In addition to the experiments in Nuf mice, we exploited this compound to study the effects of CaSR activation on Ca^2+^ homeostasis. Administration of etelcalcetide hydrochloride resulted in a dose-dependent decrease in plasma Ca^2+^ concentrations in WT mice (Figure 6A). Plasma PTH concentrations were significantly lower in mice treated with 2.5 mg/kg etelcalcetide hydrochloride compared to vehicle-treated animals (Figure 6B). Although a trend towards a reduced urinary Ca^2+^ excretion was observed in both etelcalcetide hydrochloride treatment groups compared to vehicle, this difference was not significant (Figure 6C). Nevertheless, like Nuf mice in their basal state, *Cldn14* expression was unaltered by etelcalcetide hydrochloride (Figure 6D). The renal expression of *Cyp24a1* was significantly decreased in the group that received 10 mg/kg etelcalcetide hydrochloride compared to the vehicle-treated group (Figure 6E). No significant differences were present for *Cyp27b1* (Figure 6F). There was no localization of CLDN14 to the tight junctions of the TAL for mice treated with vehicle or either 2.5 or 10 mg/kg etelcalcetide hydrochloride (Figure 6G). In the duodenum, no significant differences were present between vehicle-treated mice or etelcalcetide hydrochloride-treated mice (Figure 6H). The colonic expression of *S100g* was significantly lower in mice treated with 2.5 mg/kg etelcalcetide hydrochloride compared to the vehicle-treated group and the group treated with 10 mg/kg etelcalcetide hydrochloride (Figure 6I). In addition, the expression of *Slc8a1* was significantly increased in the colon of mice that received 10 mg/kg etelcacletide hydrochloride compared to 2.5 mg/kg etelcalcetide hydrochloride (Figure 6I).

**Figure 6.**
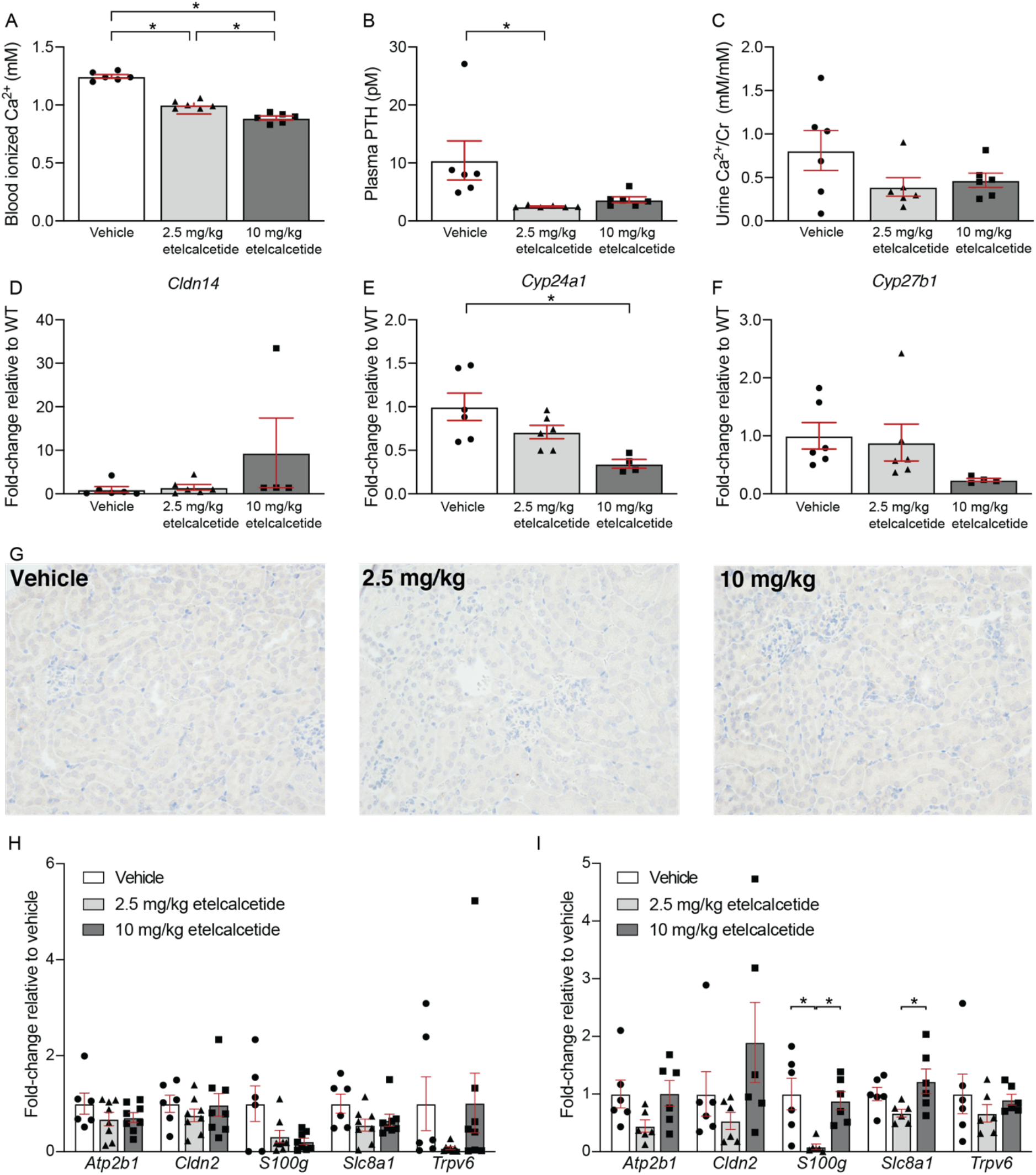
Etelcalcetide hydrochloride leads to hypocalcemia, but does not activate the renal CaSR. WT mice infused with etelcalcetide hydrochloride showed a dose-dependent reduction in blood Ca^2+^ concentrations after 7 days of treatment (*n*=6) (A). Plasma intact PTH concentrations were significantly lower in etelcalcetide treated mice than in vehicle mice (B). No statistically significant differences in urinary Ca^2+^ excretion were found between the two groups receiving etelcalcetide hydrochloride compared to vehicle-treated animals (*n*=6) (C). No changes in renal *Cldn14* were present between vehicle-treated animals and the two groups of animals that received a different dose of etelcalcetide hydrochloride (*n*=4-6) (D). The renal expression of *Cyp24a1* was significantly lower in animals having received 10 mg/kg etelcalcetide hydrochloride compared to vehicle-treated animals (*n*=4-6) (E). No significant differences were present for *Cyp27b1* between any of the groups (*n*=4-6) (F). CLDN14 was not observed localized in the tight junctions of the TAL in immunohistochemical images of mice treated with either vehicle, 2.5 mg/kg etelcalcetide hydrochloride or 10 mg/kg etelcalcetide hydrochloride (G). Etelcalcetide hydrochloride infusion to WT mice infused did not activate the intestinal CaSR, as indicated by the unaltered *Trpv6* expression in the duodenum (H) and colon (I) compared to vehicle-treated animals. The duodenal expression of other calciotropic genes was also unaltered (*n*=5-6) (H). In the colon, the expression of *S100g* was significantly reduced in mice receiving 2.5 mg/kg etelcalcetide hydrochloride compared to vehicle-and 10 mg/kg etelcalcetide hydrochloride-treated animals, while the expression of *Slc8a1* was significantly increased in animals treated with 10 mg/kg compared to 2.5 mg/kg (*n*=6) (I). Data are shown as mean ± SEM (*n*=4-6). \**P* < 0.05 by one-way ANOVA with Tukey’s *post hoc* test.

### Relationship between urinary Ca^2+^ excretion and blood Ca^2+^ concentrations in a patient with ADH1

To translate our findings to patients with ADH1, who have gain-of-function mutations in *CASR*, urinary Ca^2+^ excretion was followed over time and related to blood Ca^2+^ concentrations in a patient with ADH1. This now 17-year-old boy presented at 2 years of age with hypocalcemia and low plasma PTH levels, without hypercalciuria. This patient was followed for 15 years, resulting in an extensive collection of blood and urine samples. Sequencing of the *CASR* revealed a novel mutation (c.C2351T, p.A784V). He was treated initially with calcitriol and calcium supplementation, but had persistent hypocalcemia, sleeplessness and gastrointestinal upset, such that it was decided to start subcutaneous synthetic PTH 1-34 (3.5 units t.i.d., teriparatide; Forteo) treatment at the age of 10. He remains on this currently at a dose of 20 μg, t.i.d. Strikingly, hypercalciuria (defined as Ca^2+^/creatinine ratio > 0.6) was invariably present when he was normocalcemic (ionized blood Ca^2+^ between 1.09 and 1.25 mM) (Figure 7A). Moreover, when blood ionized Ca^2+^ concentrations were below the normal range he was less likely to have hypercalciuria (Figure 7A). Similarly, the fractional urinary excretion of Ca^2+^ was higher when blood Ca^2+^ concentrations were within the normal range, compared to most of the time when hypocalcemia was present (Figure 7B). Blood intact PTH concentrations were strongly suppressed regardless of blood Ca^2+^ levels (Figure 7C). Unfortunately, he developed nephrocalcinosis at 9 years of age (Figure 7D). His renal function remains preserved with a plasma creatinine of 69 μmol/L at last follow-up.

**Figure 7.**
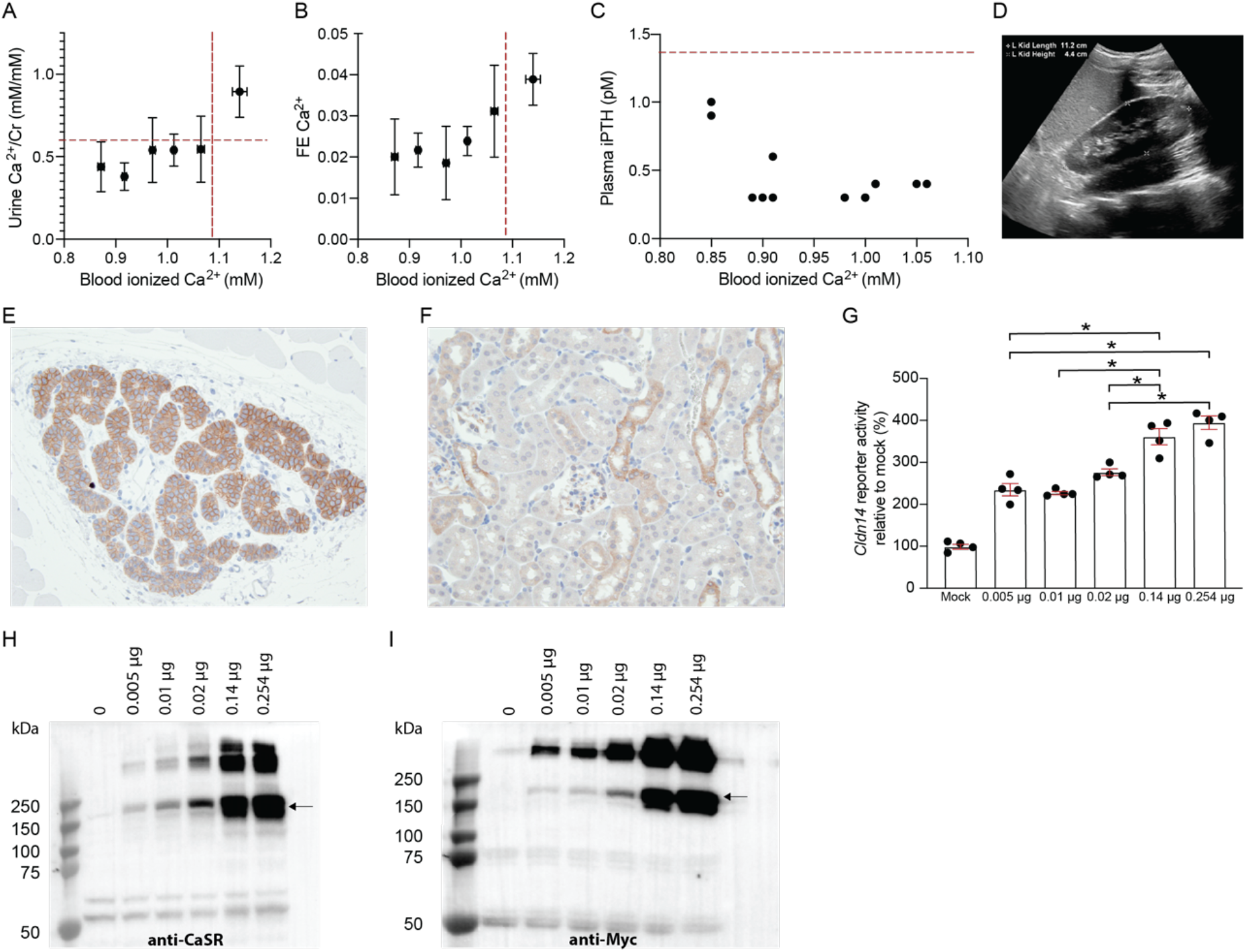
Ca^2+^ homeostasis in a patient with ADH1 and relationship between CaSR abundance and CaSR activity. Treatment of a patient with ADH1 in order to normalize blood Ca^2+^ levels were associated with the occurrence of hypercalciuria when the patient was normocalcemic (*n*=6-19 individual measurements per cluster) (A). In addition, fractional urinary excretion of Ca^2+^ (FE Ca^2+^) was increased after treatment leading to normocalcemia (*n*=6-19 individual measurements per cluster) (B). Plasma intact PTH levels (iPTH) were suppressed during all collection periods (*n*=6-19 individual measurements per cluster) (C). Medullary calcinosis, a complication of treatment-induced hypercalciuria in ADH1, was observed by ultrasound (D). Immunohistochemistry revealed a marked immunoreactivity of CaSR in the parathyroid (E). CaSR immunoreactivity was also detected in the thick ascending limb of Henle’s loop in the kidney, albeit at a lower intensity (F). After transfection of HEK293 cells with different amounts of a construct encoding the human CaSR (hCaSR) and a fixed amount of a CLDN14-promoter reporter construct, higher reporter signal was obtained after transfection of cells with a higher amount of construct (*n*=4 independent experiments) (G). The increased hCaSR expression was confirmed by western blotting with an anti-CaSR antibody (H) and with an anti-C-myc antibody (I). Representative immunohistochemical images and western blots are shown; arrows indicate dimeric CaSR signal. Data are shown as mean ± SEM. \**P* < 0.05 by one-way ANOVA with Tukey’s *post hoc* test.

### In vitro activation of the CaSR

Data from the Human Protein Atlas suggests that the CaSR is expressed higher in the parathyroid than in peripheral tissues, which could help explain the different CaSR activation thresholds between the parathyroid and kidney (30). Staining of the CaSR in mouse indicated higher cellular abundance of the CaSR in the parathyroid (Figure 7E) compared to the kidney TAL segments in the cortex (Figure 7F). To test the effect of increasing CaSR expression in the presence of a consistent extracellular Ca^2+^ concentration sufficient to activate the CaSR (1.5 mM Ca^2+^), a CaSR-responsive *Cldn14* promoter reporter was transfected with increasing amounts of a myc-tagged CaSR into HEK293 cells. Luminescence (and thus CaSR activity) significantly increased proportionately with increasing amounts of the CaSR construct transfected (Figure 7G). To confirm that increased plasmid transfection resulted in increased protein expression, we immunoblotted for CaSR expression after transfection with either an anti-CaSR antibody (Figure 7H) or anti-myc antibody (Figure 7I). The indicated bands showed monomeric and dimeric CaSR expression (31). These results illustrate that at an extracellular Ca^2+^ concentration of 1.5 mM, additional receptor activity can be achieved by increasing the abundance of the receptor.

## Discussion

The CaSR in the parathyroid and peripheral CaSRs play an important role in maintaining systemic Ca^2+^ balance by defending against hypercalcemia (9). In the kidney, activation of the CaSR strongly increases *Cldn14* expression, leading to the localization of CLDN14 to the tight junctions of the TAL. This reduces the permeability of the tight junction and thereby increases urinary Ca^2+^ excretion (7, 8). Using *Cldn14* as a surrogate marker of CaSR activity, we examined renal CaSR activation in a mouse model of ADH1 and following pharmacological CaSR activation. Our data strongly suggest that activation of the renal CaSR only occurs when circulating Ca^2+^ concentrations reach sufficiently high concentrations, even though the receptor harbors overactivating mutations or is activated pharmacologically, because activation of the CaSR in the parathyroid reduces circulating Ca^2+^ concentrations. This is based on the following observations: *i)* the fractional excretion of Ca^2+^ and furosemide sensitivity were unaltered in Nuf mice *ii)* CaSR-dependent increases in the expression of *Cldn14* in the TAL were absent in Nuf mice or following administration of the calcimimetic etelcalcetide hydrochloride. *iii)* CLDN14 expression only increases following elevations in plasma Ca^2+^ concentrations above baseline levels in both WT and Nuf mice. i*v)* A patient with ADH1 displays persistently suppressed blood intact PTH levels under hypocalcemic conditions, while hypercalciuria mainly occurred when treatment succeeded in normalizing blood ionized Ca^2+^ levels.

The underlying mechanism mediating a differential response of the parathyroid CaSR versus peripheral CaSRs at a given plasma Ca^2+^ concentration remains to be elucidated. Data from the Human Protein Atlas suggests that CaSR is highest expressed in the parathyroid compared to other tissues (30). Accordingly, CaSR abundance appeared higher in mouse parathyroid than kidney. Therefore, pharmacological activation or gain-of-function mutations in the CaSR may lead to greater total activation of the CaSR in the parathyroid at the same plasma Ca^2+^ levels. Furthermore, *in vitro* transcription of *Cldn14* was increased after transfection with increased amounts of CaSR-encoding construct. Tissue-specific signaling pathways may also facilitate differential CaSR activation. In fact, CaSR-dependent signaling in the TAL may differ substantially from that reported in the parathyroid (19).

Based on the differential activation of the parathyroid CaSR and the CaSRs in peripheral tissues observed here, we propose that CaSR activation in the parathyroid and subsequent modulation of PTH secretion is vital for maintaining normocalcemia. Importantly, when *Cldn14* expression is plotted against blood ionized Ca^2+^ concentrations for all WT and Nuf/+ mice in the Nuf experiments, curve fitting using a power trend line resulted in a similar relationship for WT (R^2^ = 0.44) and Nuf/+ mice (R^2^ = 0.46) (Figure 8A). However, the relationship is shifted leftwards for Nuf/+ mice as would be expected with increased CaSR sensitivity. This indicates that although the renal CaSR is inactive under baseline blood Ca^2+^ concentrations in Nuf/+ mice, it becomes activated at a lower blood Ca^2+^ concentration than WT mice. Furthermore, a similar relationship is present for both genotypes between plasma PTH concentrations and blood ionized Ca^2+^ concentrations (R^2^ = 0.82 and 0.78 for WT and Nuf/+, respectively) (Figure 8B). However, plasma PTH were already decreased in Nuf/+ mice at baseline blood Ca^2+^ levels, indicating that the parathyroid CaSR is already activated in Nuf/+ mice at baseline and supports the notion of differential activation thresholds in parathyroid and kidney.

**Figure 8.**
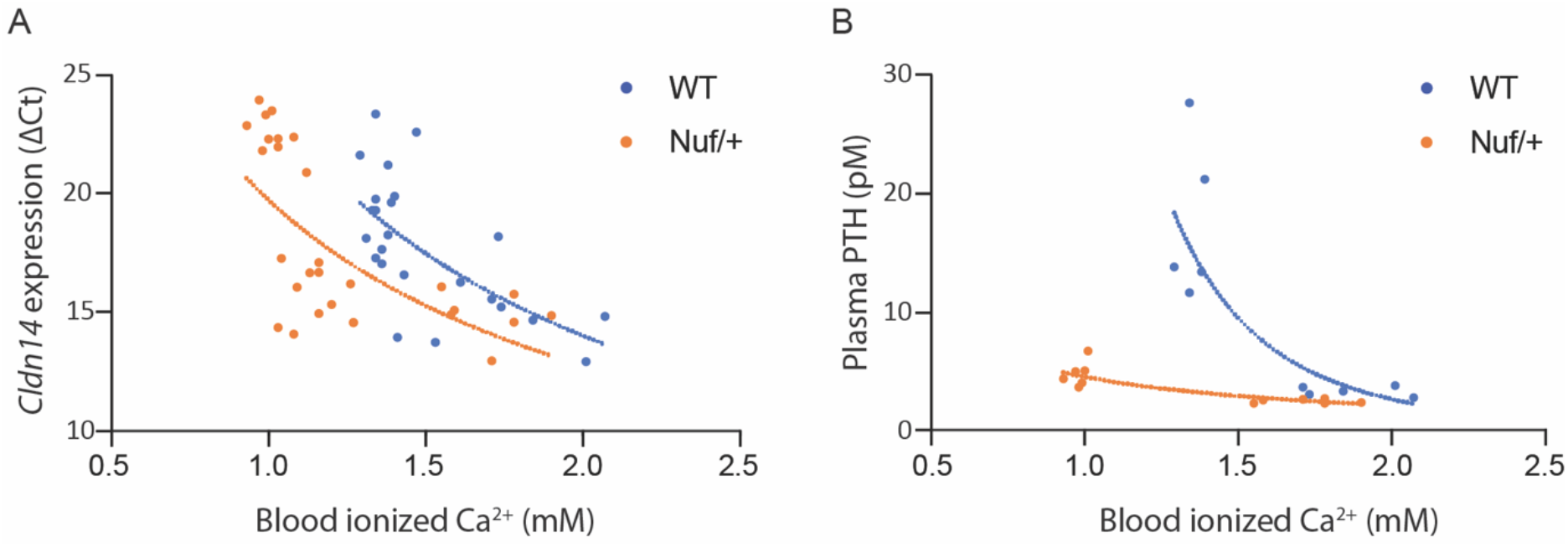
Relationship between blood Ca^2+^ levels and renal *Cldn14* expression or plasma PTH concentrations in Nuf/+ mice. Pooling blood Ca^2+^ concentrations and *Cldn14* gene expression (ΔCt; *Cldn14* Ct – *18S* Ct) data from all experiments revealed a non-linear relationship, which could be described with a power trend line that fitted the data points equally well for both genotypes (R^2^ = 0.44 for WT and 0.46 for Nuf/+ mice). A leftward shift was observed for Nuf/+ mice (A). A similar non-linear relationship was present between plasma PTH concentrations and blood Ca^2+^ concentrations (R^2^ = 0.82 for WT and 0.78 for Nuf/+), yet PTH was already markedly suppressed in the Nuf/+ mice.

In contrast, the activation of peripheral CaSRs, including the renal and intestinal CaSR, might be limited to states of hypercalcemia where a reduction in PTH fails to reduce plasma Ca^2+^ levels. Accordingly, a previous study showed that *Pth*^-/-^ mice, but not *Casr*^-/-^ *Pth*^-/-^ double knockout mice were still able to withstand hypercalcemic challenges (32).

The differential activation of CaSRs at a given blood Ca^2+^ concentration in the parathyroid *vs.* kidney and intestine is in accordance with the phenotype of patients with ADH1. These patients exhibit hypocalcemia and reduced plasma PTH concentrations, reflecting activation of the parathyroid CaSR (11). However, untreated ADH1 patients are hypocalciuric and often display hypercalciuria after treatment (11–15). In the current study, hypercalciuria also occurred when blood Ca^2+^ levels were increased in Nuf mice, suggesting that the renal CaSR is initially inactive in patients with ADH1 and only becomes activated after increased plasma Ca^2+^ concentrations. However, this may also depend on how much a given mutation activates the receptor, such that some ADH1 patients may have renal receptor activation at lower plasma Ca^2+^ concentrations, even though others will not. In fact, gain-of-function mutations in *CASR* can also cause ADH1 with a Bartter-like syndrome, including hypercalciuria before Ca^2+^ or vitamin D supplementation (33–35). Accordingly, two other mouse models with different gain-of-function mutations in either the transmembrane or extracellular domain of CaSR, displayed a clear increase in urinary Ca^2+^ excretion at baseline, despite having similar blood Ca^2+^ concentrations to the Nuf mice (27). Both of these hypercalciuric mouse models are based on human ADH1 mutations, while one mutation was also found in an ADH1 patient with a Bartter-like syndrome (27, 35, 36). These differences illustrate that the phenotype resulting from genetic overactivation of the CaSR, can be more severe depending on the exact mutation and hence the level of CaSR activation at a given plasma Ca^2+^ concentration.

In line with this, etelcalcetide hydrochloride-treated mice are hypocalcemic without activation of the renal CaSRs. In contrast, another calcimimetic, cinacalcet, potently activates the CaSR and increases CLDN14 expression and urinary Ca^2+^ excretion (7). Like the differential phenotype of parathyroid and renal CaSR activation in ADH1, this shows that more potent modulation of the CaSR can allow activation of the renal CaSR in the presence of lower circulating Ca^2+^ concentrations.

Importantly, Nuf mice have a missense mutation in the fourth transmembrane domain of CaSR(18), while transmembrane mutations in human ADH1 patients, including the one described here, are usually present in transmembrane domains 5 and 6 (11). Nevertheless, the observation that both Nuf mice and the ADH1 patient developed hypercalciuria after normalization of plasma Ca^2+^ concentrations suggests that the Nuf mice are an adequate model of ADH1.

Our results have clear implications for the management of patients with ADH1. Hypercalciuria is a significant risk factor for nephrocalcinosis and consequently renal dysfunction. Based on our findings, treatment-induced hypercalciuria in ADH1 can be avoided by titrating the plasma Ca^2+^ level to one just below the normal range. This prevents the activation of the renal CaSR and thus hypercalciuria while hopefully preventing symptoms of hypocalcemia. In the aforementioned patients as well as in a subset of patients with more potent activating CaSR mutations and hence larger renal responses at even smaller increments in plasma Ca^2+^ or at any given plasma Ca^2+^ contraction, potential specific treatment for ADH1 could be achieved by combining CaSR antagonists with vitamin D treatment to raise plasma Ca^2+^, while limiting urinary Ca^2+^ wasting. This strategy would reduce the increased risk of developing stone disease, calcifications and kidney disease while at the same time raising plasma Ca^2+^ levels to remain asymptomatic.

In conclusion, we show here that the peripheral CaSRs are differentially activated from the parathyroid. These findings could explain why hypercalciuria is absent in many patients with ADH1 and only occurs after treatment of hypocalcemia and aid in the clinical management of these patients.

## Author contributions

HD and RTA conception; WvM, HD and RTA designed research; WvM, RT, RTA and HD performed experiments; WvM, RT, RTA and HD analyzed data; WvM, RT, RTA and HD interpreted results of experiments; WvM prepared figures; WvM and HD drafted the manuscript; WvM, RT, RTA and HD edited, revised and approved the final version of the manuscript.

## Acknowledgments

The authors thank Inger Nissen, Lene Bundgaard Andersen, Mohamed Abdullahi Ahmed, Rasmus Andersen and Kenneth Andersen at the University of Southern Denmark and Wanling Pan at the University of Alberta for expert technical assistance. This work was supported by grants from Erasmus+ 2018/E+/4458087 awarded to WH van Megen, the Canadian Institutes for Health research to RT Alexander and the Novo Nordisk Foundation, the Beckett Foundation, the Carlsberg Foundation, the Lundbeck Foundation and Independent Research Fund Denmark awarded to H. Dimke. Dr. R. Todd Alexander is the Canada Research Chair in Renal Tubular Epithelial Transport Physiology and is a Distinguished Researcher of the Stollery Children’s Hospital Foundation.

## Disclosures

The authors have declared that no conflict of interest exists.

---

**Table S1.**
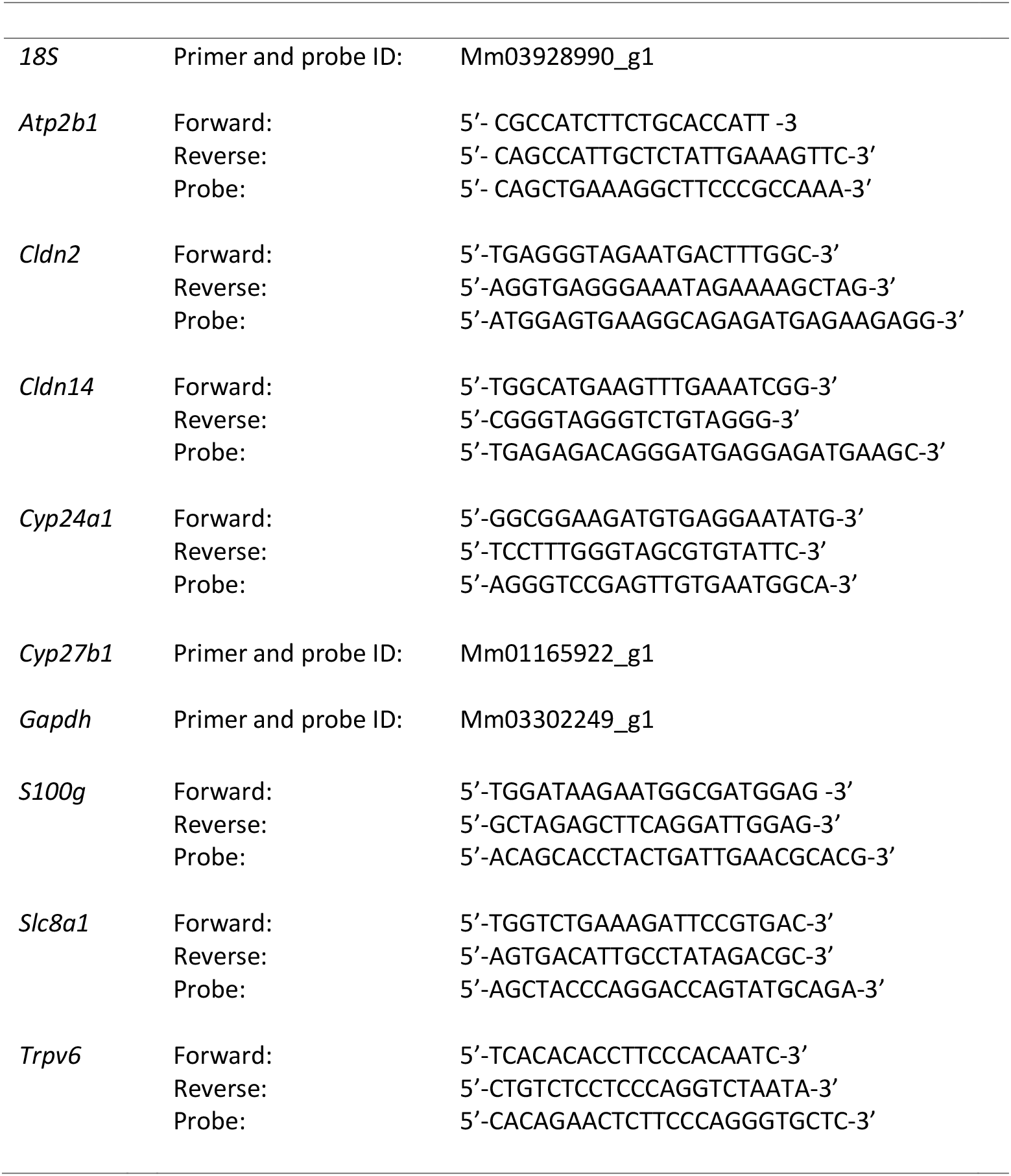
Real-time PCR primers and probes.

## Notes

### Competing Interest Statement

The authors have declared no competing interest.

